# Modeling one-shot interceptions in fruit-catching fish

**DOI:** 10.64898/2026.09.17.752477

**Authors:** Aria Azari-Pour, L. Mahadevan

## Abstract

Interception of moving objects is a universal pattern of animal behavior with many industrial applications. Inspired by the behavior of the characid fish *Brycon guatemalensis*, which catches fruits falling onto the surface of a river, we study one-shot (i.e., open-loop) interceptions using both stochastic optimal control theory and reinforcement learning. In the first instance, an agent actuates a control (wait time before launch and heading angle) to meet its target, given a noisy estimate of the target initial state. In a minimal setting, where launch velocity and actuated control are constants, we show that the optimal wait time is attenuated by noise in both the target measurement and control actuation, as well as agent energy conservation. We then derive an upper bound for the heading angle variance which scales as the ratio of the non-vanishing optimal interception error to the initial planar separation. Complementing our control framework, our deep reinforcement learning approach shows that the neural network representing the fish first learns the value function for the interception, then learns how to aim, and finally learns the optimal wait time. Beyond the specific problem at hand, our results may also have applications to problems with little time for feedback or corrections such as perching, landing, and docking.

## I. INTRODUCTION

Interception is an essential behavioral paradigm observed across the animal kingdom in which an animal intercepts prey and other objects [1–4]. In humans, interception mechanisms have been particularly well-studied [5–8], especially in ball-catching team sports such as soccer, rugby, and basketball where the term *interception* colloquially refers to catching a pass intended for a member of the opposing team. The topic has also recently become relevant in artificial settings such as robotics [9–12], with a dramatic recent example being that of a robot beating an expert human in table-tennis [13] by repeatedly and robustly intercepting a moving table-tennis ball with a paddle.

From a mathematical perspective, an interception can be framed as an optimal control problem in which an agent controls some trajectory of interest, often its own, to coincide with a separate target trajectory at some point in time [14]. Interceptive mechanics are classified based on the control loop of the agent which depends on the timescale of target dynamics. *Adaptive* (closed-loop) interceptions are those where the timescale of target dynamics is much larger than the timescale of processing and action in the agent, allowing for feedback and corrections to initial control inputs. The control process is adapted to a filtration which represents the continuous tracking of a stochastic process modeling the target dynamics [15]. *One-shot* (open-loop) interceptions, on the other hand, are those where the target dynamics occur over a timescale at most as large as the timescale of processing and action in the agent. As a result, there is an initial decision and control actuation, after which the control input remains constant during the target dynamics.

While there is extensive work on adaptive interceptions and filtered optimal control [16], there is much less work on one-shot interceptions despite its ubiquity in everyday situations [17]. A familiar example of a one-shot interception is quickly catching a heavy object falling off the edge of a table, when there is barely enough time to register motion before an action is attempted. Several other examples are shown schematically in Fig. 1, from baseball catchers receiving fastballs to archers on horseback [18].

**FIG 1:**
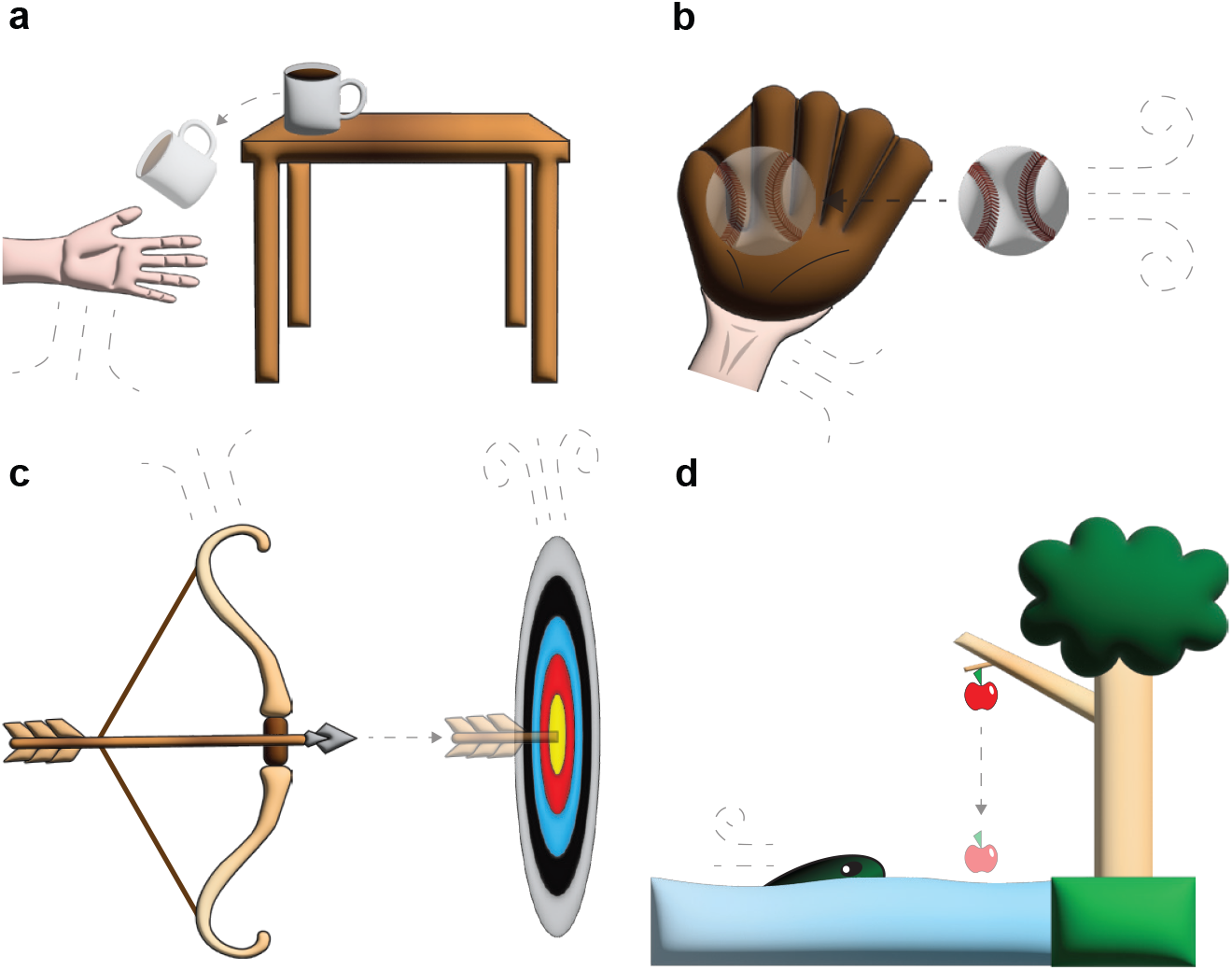
Archetypal examples of one-shot (open-loop) interceptions. (a) Catching an object falling off the edge of a table. (b) A baseball player catching a baseball moving very quickly. (c) A moving archer firing an arrow at a nearby dynamic target. (d) *Brycon guatemalensis* intercepting a fruit falling from a tree.

A prototypical one-shot interception is observed in the behavior of the Central American characid fish *Brycon guatemalensis*, which intercepts fruits falling onto the surface of the water on which it swims [19–21]. *B. guatemalensis* (hereafter referred to as *fish* throughout this paper) is a frugivore whose adult diet consists mostly of leaves and figs from the riparian tree *Ficus insipida*. When a fig fruit from *F. insipida* falls from above the water, submerged *B. guatemalensis* wait near the surface before initiating a *C* -start mechanism to propel themselves towards the impact point of the fruit. The *C* -start is a rapid, accelerating, reflexive movement seen in many larval fish [22, 23] which leads to a burst of movement to escape predation. It has been proposed that the *C* -start has been repurposed in some adult fish, such as *B. guatemalensis*, as a behavioral strategy for feeding [24]. In addition to the question of optimal control, there is an equally important question regarding the role of agent learning in one-shot interceptions. Reinforcement learning (RL) methods [25, 26] are a useful tool to model learning strategies where an agent must interact with the environment through trial-and-error. RL has been shown as a mechanism for learning in both songbirds and humans [27, 28].

In this paper we investigate both the control and learning aspects of one-shot interceptions. In Sec. II, we describe the general optimal control model of one-shot interceptions. Then, in Sec. III, we apply the optimal control model to the fruit-fish interception. In Sec. IV, we solve the stochastic optimal control problem for the fruit-fish interception with noise and in Sec. V, the results are compared with experimental observations from Ref. [21]. The role of learning in the fruit-fish interception is explored in Sec. VI, where we describe a deep reinforcement learning architecture to model learning of the optimal policies of one-shot interceptions. The results of the reinforcement learning model are described in Sec. VII for one-shot interceptions in the noiseless limit.

## II. GENERAL ONE-SHOT INTERCEPTION MODEL

A one-shot interception occurs between a dynamic *target* and a dynamic *agent* attempting to intercept the target. Let **x**(*t*) = (*x*_1_(*t*), *x*_2_(*t*), *x*_3_(*t*)) be the trajectory of the target and **y**(*t*) = (*y*_1_(*t*), *y*_2_(*t*), 0) the trajectory of the agent in standard Euclidean coordinates. The target dynamics occurs over a time horizon [0, *T* ]. We assume that the agent is initially at rest and is confined to move in the plane *z* = 0 onto which the target will land at *T*, as well as initially *x*_3_(0) *>* 0 and the target is accelerated in the ( *−z*)-direction by gravity with acceleration *g*.

Following the motion of the target at *t* = 0, the agent decides to wait for a fraction *w* of *T*, during which time it aligns itself at a heading angle Φ in the plane *z* = 0. At *t* = *wT*, the agent instantaneously accelerates to a velocity *v* and moves in the direction of the heading angle Φ in the plane *z* = 0 towards the *impact point* of the target, **x**(*T*). The *impact time* for the target to reach the plane *z* = 0 is *T* = (2*x*_3_(0)*/g*)^1*/*2^. We denote by **u** = (*w*, Φ) the control actuated by the agent at *t* = 0.

The dynamics of a one-shot interception is entirely determined by the egocentric relative coordinates

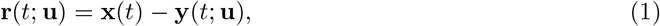

for *t ∈* [0, *T* ], where we emphasize that the control **u** affects only the agent dynamics. For fixed target dynamics **x**(*t*), the relative coordinates are determined by the initial choice of **u** at *t* = 0.

Define

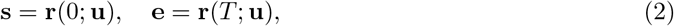

as the initial state and terminal error, respectively. For simplicity, we fix **y**(0; **u**) = **0** such that the initial state **s** = **x**(0) is independent of the control. If **s** is fixed, then the terminal error has the functional form **e** = **e**(**s**, *v*, **u**). The initial state **s** and the control **u** are assumed to be random variables.

The agent aims to minimize **e** in order to intercept the target subject to some physically limiting constraints. For instance, the finite speed *v* implies that it cannot reach targets that are too far away in a finite time. In a one-shot interception, *v* and **u** remain constant over [0, *T* ] as there is not enough time for dissipative effects to dampen the initial launch velocity, nor for the agent to correct the initial control actuation. The assumption of constant velocity over *T* is equivalent to the condition that drag effects are small on timescales associated with the *C* -start [29, 30].

Formally, we may cast a one-shot interception in terms of an an optimal control problem as follows: *determine the optimal control* **u**^∗^ *that consists of (a constant) optimal wait fraction w*^∗^ *and (a constant) optimal heading angle* Φ^∗^, *to be actuated by the agent as* argmin

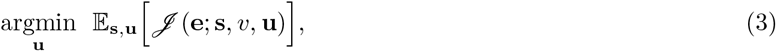

where

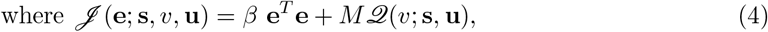

*subject to the constraints*: the speed *v* is constant on [0, *T* ], *x*_3_(*T*) = 0, *x*_3_(0) *≥ g*∥**x**(*T*) ∥^2^*/*(2*v*^2^), *y*_3_(*t*) = 0, *w∈* [0, 1), Φ *∈* [*−π, π*).

The first term on the right-hand side of Eq. (4) is the squared-error of the interception with *β >* 0 the *hunger* parameter that encourages interception. Within the second term, *ℒ*(*v*; **s, u**) is an inertial penalty for interceptions (to be defined by the context) which represents fatigue through kinetic energy expenditure in biological interceptions or electrical energy usage in robotics systems and *M >* 0 is the *inertia* (or *fatigue*) parameter that penalizes energy expenditure^1^.

The agent must measure the initial state to actuate the optimal control and the control actuation itself contains noise through signal transduction and motor imprecision. Then, the expectation in Eq. (3) is taken with respect to the distributions of the initial state and control. Throughout this paper, expectations will be assumed to be taken with respect to the distribution of a random variable given by the context.

## III. ONE-SHOT INTERCEPTIONS IN FRUIT-CATCHING FISH

A one-shot interception is conducted by a fish confined to swim on the surface of a river, given by the plane *z* = 0, onto which a fruit is falling. As shown in Fig. 2, the fruit falls from rest at *t* = 0 under the force of gravity with trajectory 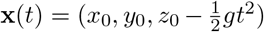. The trajectory of the fish is **y**(*t*; **u**) = *vW*(*t*)(cos Φ, sin Φ, 0), where

**FIG 2:**
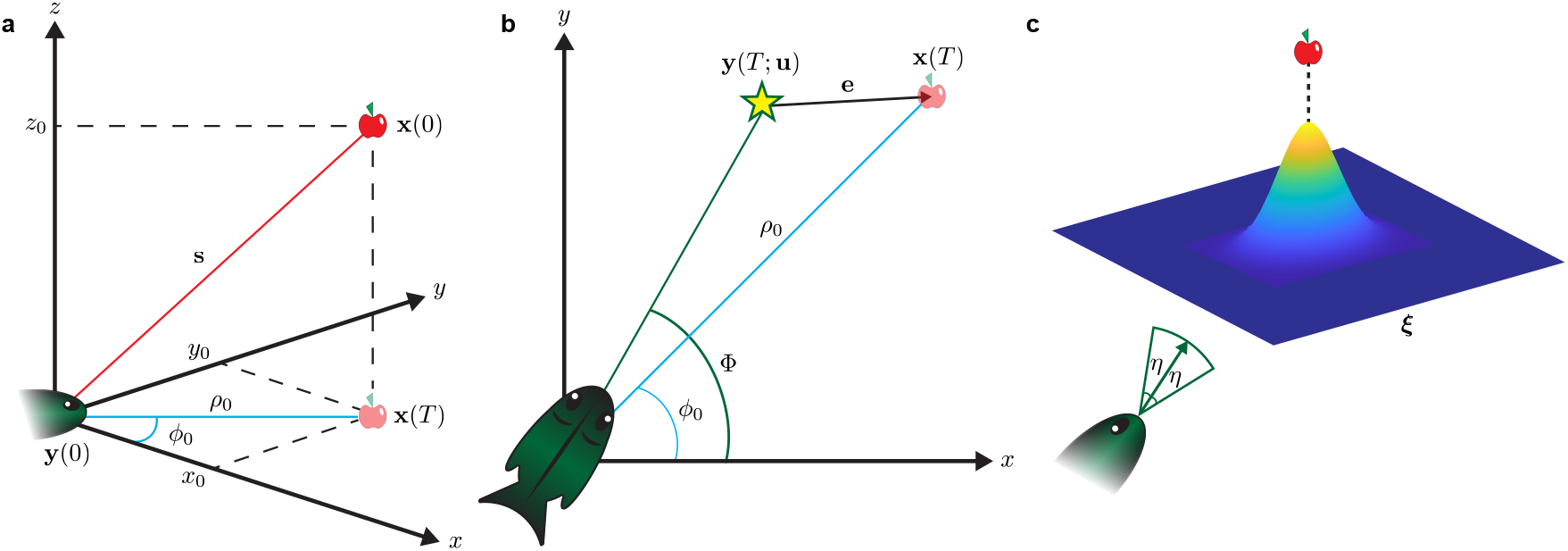
Fruit-catching fish as a prototypical model of one-shot interceptions. (a) A fruit falls under the force of gravity. The egocentric coordinate system is shown with initial state **s** = **r**(0; **u**). (b) At *t* = 0, the fish decides to wait for a fraction *w* of the impact time *T* of the fruit, during which time it aligns itself at a heading angle Φ towards the impact point **x**(*T*). Then, at *wT*, the fish swims at a constant speed *v* in the direction of the heading angle Φ. The interception error **e** is the planar distance between the fish and the fruit at *T* . (c) There is noise in both the measurement of the impact point **x**(*T*) = (*x*_0_, *y*_0_, 0) by the fish and the heading angle Φ, given by ***ξ*** and *η*, respectively.

**FIG 3:**
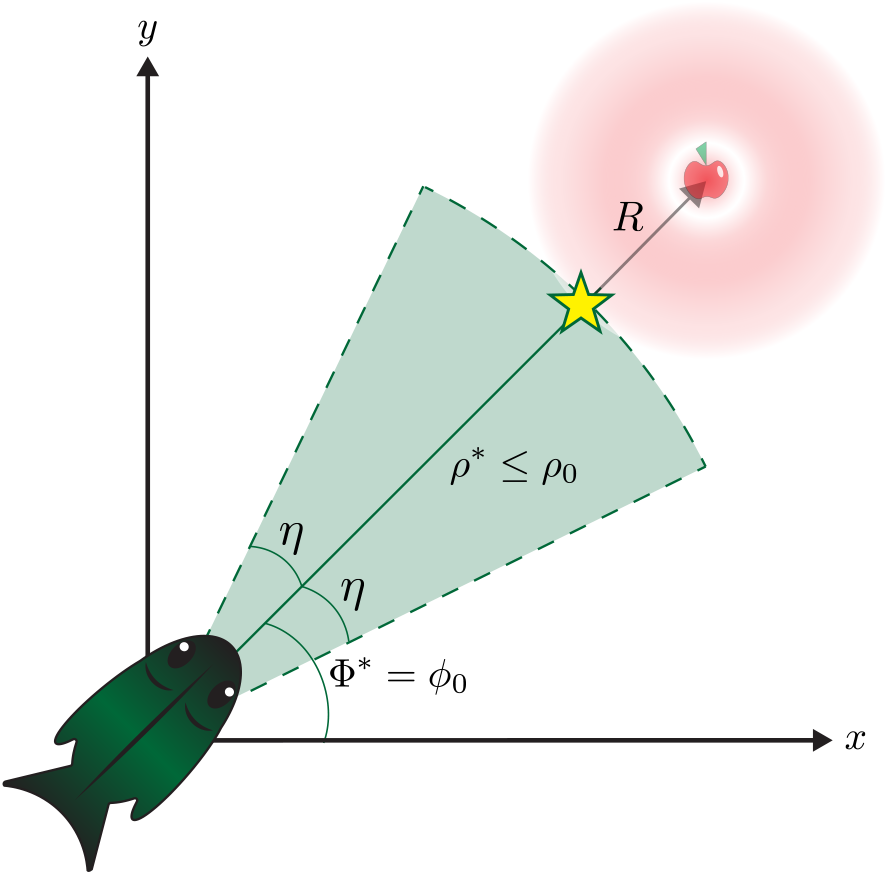
Stochastic optimal control for one-shot interceptions. The fish actuates the optimal control **u**^∗^ = (*w*^∗^, Φ^∗^) with *ρ*^∗^ = (1 *− w*^∗^)*vT ≤ ρ*_0_ the optimal swim distance. The expressions for Φ^∗^ and *w*^∗^ are given in Eqs. (12) and (14), respectively. The optimal error has magnitude *R* = ∥**e**^∗^∥ = *ρ*_0_ − *ρ*^∗^ *≥* 0.

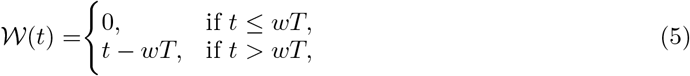

is the cumulative time the fish has been swimming.

The relative position of the fruit with respect to the fish is

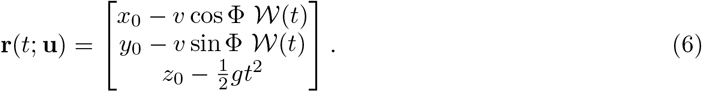

The interception has initial state **s** = (*x*_0_, *y*_0_, *z*_0_) and terminal error

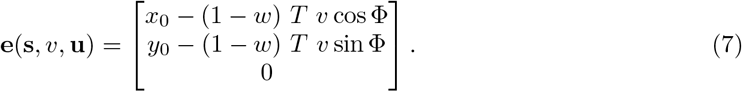

In order for the fish to be able to intercept the fruit, the impact time must be at least as large as the time for the fish to swim to the impact point, *T ≥ ρ*_0_*/v*, with 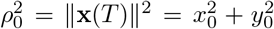. As *v* is constant during the interception, then one may consider lengths as a timescale with *v* the conversion factor. The geometric constraint on the initial state **s** is

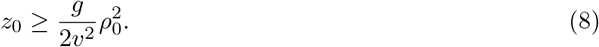

If Eq. (8) does not hold, it is physically impossible for the fish to reach the impact point by the time the fruit has fallen on the surface of the water.

## IV. STOCHASTIC OPTIMAL CONTROL OF ONE-SHOT INTERCEPTIONS

The fish measures the initial state **s**, and consequently the impact point **x**(*T*), with imprecision. Let **s** = E[**s**] +***ξ*** be the measurement of the initial state by the fish where ***ξ*** = (*ξ*_1_, *ξ*_2_, 0) is a vector of zero-mean random noise with variance 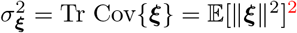. Also, actuation of the heading angle by the fish is imprecise. The heading angle is Φ = E[Φ] +*η* where *η* is zero-mean random noise with variance 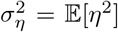. We consider an inertial term of the form *ℒ*(*v*; **s, u**) = *v*^2^(1 *−w*)*T/*2^3^. The objective function in Eq. (4) may be written explicitly as

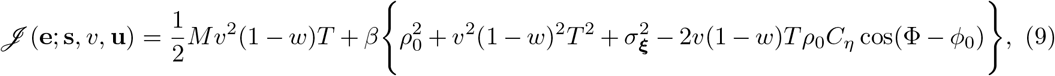

where *C*_*η*_ = E[cos *η*] *≤* 1 is an attenuation factor from the angular noise.

The optimal control **u**^∗^ = (*w*^∗^, Φ^∗^) requires that the gradient with respect to the control vanishes, *∇*_**u**_*J* (**u**^∗^) = **0**, which yields

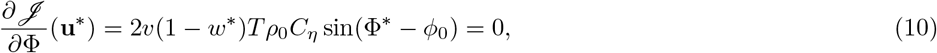

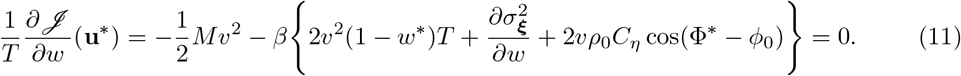

The optimal heading angle Φ^∗^ is then given by solving Eq. (10), which gives

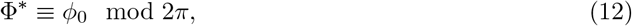

*?*where we assume *w*^∗^ ≠ 1 for an interception with finite swim speed. Noise does not affect optimality of the heading angle and the fish should always aim at the impact point of the fruit irrespective of noise.

The optimal wait fraction *w*^∗^ is found by solving Eq. (11). Assume the impact point variance 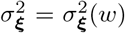 is a monotone-decreasing smooth function of *w*. As the fish waits longer, the impact point variance decreases and the estimate of the impact point in the initial state **s** becomes more accurate as the fruit is closer to the fish. We consider 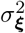 to linear order in *w* and define

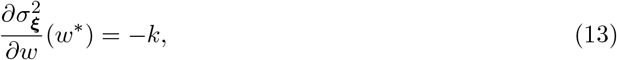

where *k >* 0 is the (constant) *variance decay rate* at *w*^∗^. The negative sign emerges due to the increase in measurement accuracy by the fish with decreasing relative position of the fruit. Solving Eq. (11) at the optimal heading angle, we have

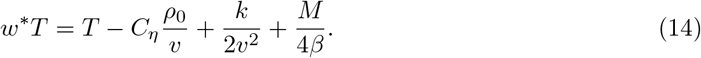

The optimal swim time is (1 *− w*^∗^)*T*, which yields an optimal distance traveled by the fish,

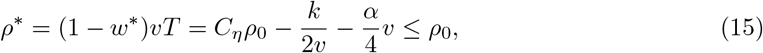

where *α* = *M / β* is the *laziness* of the fish. The inequality in Eq. (15) follows as *C*_*η*_ *≤* 1 and *k, v, M, β ≥* 0.

The optimal error vector **e**^∗^ has direction given by the optimal heading angle *ϕ*_0_ and magnitude *R* = *ρ*_0_ *− ρ*^∗^ *≥* 0, which we define as the *interception radius*

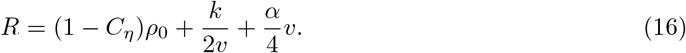

We may write the angular attenuation factor as 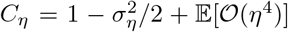. For small angular noise *η*,

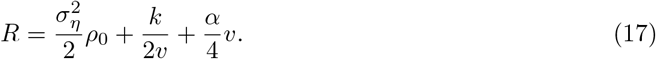

we have The heading angle variance is then

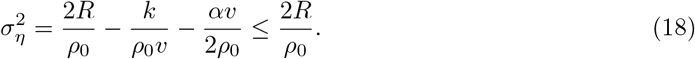

The surprising result that *R ≥* 0 in stochastic interceptions is due to the following factors. First, we have assumed that 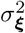 is a monotone decreasing function of *w*, such that the impact point measurement becomes more accurate as the agent waits longer and the target becomes closer to the agent. There is a competition between the agent traveling at the wait time for which *R* = 0 and waiting longer to increase the measurement accuracy of the impact point, such that *R >* 0. Second, heading angle noise also results in a lateral deviation from the direction given by Φ = *ϕ*_0_. Attenuating the swim distance then mitigates this lateral deviation such that the stochastic squared-error is minimized for an interception. Finally, the interplay of the inertial term and the squared-error term, and consequently the magnitude of *α*, affects the optimal wait time. A lazy agent will wait longer with *R >* 0, not for minimizing the expected squared-error of the interception, though for minimizing the swim time for energy preservation.

## V. EXPERIMENTAL FRUIT-CATCHING FISH INTERCEPTIONS

We now turn to connect our results to the experiments [21] on the interceptions of *B. guatemalensis* shown in Fig. 4. When fruit are dropped onto the surface of the water on which the fish swim, the school of fish perform a *C* -start mechanism and swim towards the impact point of the fruit (Fig. 4a). The interception occurs over a timescale of *T ≃* 1 second, consistent with the timescale of a one-shot interception. Interestingly, none of the fish in the experiments actually arrived at the impact point when the fruit landed on the surface of the water, matching the prediction of a non-vanishing interception radius *R*, labeled for the fruit-fish interception in Fig. 4a. Moreover, there is significant variability in the swim distances and heading angles of the fish during the interception, which may be explained by the variability of laziness parameters *α* for the school of fish, as well as stochastic realizations of measurement and control actuation noise.

**FIG 4:**
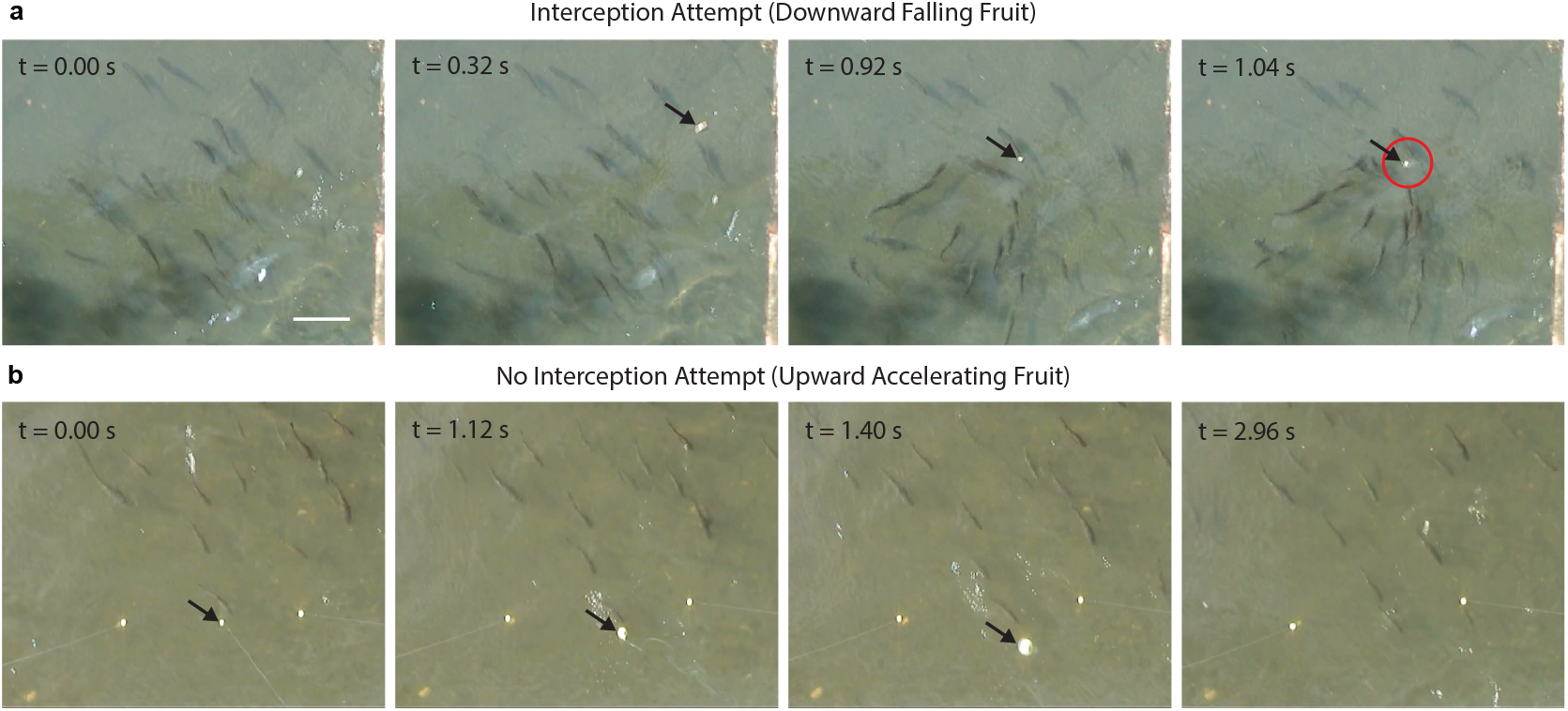
Experimental one-shot interceptions of the Central American characid fish *Brycon guatemalensis*. (a) A fruit (black arrow) is dropped above a school of *B. guatemalensis*. The fish wait for some time and then begin swimming towards the fruit. The fruit lands on the surface of the water at time *T* = 1.04 s and the interception radius *R* is the radius of the red circle. (b) Three fruits are initially below the surface of the water and attached to wires. The middle fruit (black arrow) is artificially accelerated upwards using the wire. Time *t* is measured in seconds (*s*). Scale bar = 50 cm. Figure adapted with permission from Ref. [21].

Additionally, experiments in [21] tested the interceptions when fruit were artificially accelerated upwards with an acceleration of *g* from underneath the surface of the water (Fig. 4b). The fish did not swim towards the fruit in this situation, either underneath or above the surface of the water. In the model presented in this paper, a fruit underneath the surface of the water would have *z*_0_ *<* 0 and therefore the initial geometry would not satisfy Eq. (8), at which point the fish would not attempt the interception. If *g <* 0, then the brain of the fish computes that *T* is not a real number and the fish do not attempt the interception. It follows that it is necessary to have both *z*_0_ *>* 0 and *g >* 0 for the fish to attempt the interception.

The precision of the *C* -start mechanism in fruit-catching fish may be investigated using Eq. (18). In Fig. 4, the values of *R* = 25 cm, *ρ*_0_ = 40 cm are estimated for one of the fish. Therefore, 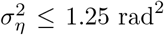 . During an interception, there is a lateral deflection *ηρ*_0_ induced by the heading angle noise *η*. We assume that the fish has an intrinsic tolerance *ε >* 0 for this deflection such that an interception is unsuccessful if *η > ε/ρ*_0_. The probability that a fish misses the fruit during an optimal interception, given a tolerance *ε*, is

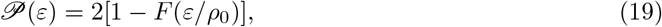

where *F* (*η*) is the cumulative distribution function of *η*. The factor of two in Eq. (19) arises from the two conditions *η > ε/ρ*_0_ and *η < −ε/ρ*_0_, where we assume the probability distribution of *η* is symmetric about *η* = 0.

If we assume *η* is a von Mises (circular normal) random variable, then *η* has distribution *p*(*η*) = exp(*κ* cos *η*)*/*(2*πI*_0_(*κ*)), with *I*_*n*_(*κ*) the *n*th-order Bessel function of the first kind and *κ* the concentration such that 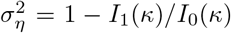. Then if *ε* = *ρ*_0_*/*10, one has *P*(*ε*) *≃* 0.90. Such a large probability for missing the fruit during an interception is observed in the experimental interceptions [21]. Surprisingly, the result suggests that the *C* -start is inefficient. However, the act of feeding may not rely on the one-shot interception itself. Our result predicts that the one-shot interception of the fruit-fish may serve to allow the fish to arrive near the fruit without feeding, and then use a separate mechanism to acquire the fruit for feeding. In this sense, the fruit-fish *C* -start mechanism and one-shot interception serve as a competitive strategy to acquire falling fruits in a school of fish, instead of a purely behavior strategy for feeding.

## VI. AGENT LEARNING IN ONE-SHOT INTERCEPTIONS

We now switch from a control theoretic perspective to consider the fish as a prototypical example of learning and use a deep RL approach [32, 33] to probe the process of learning in the fish which may be exported for a robotic controller. The architecture of the neural network for RL uses the actorcritic paradigm, as shown in Fig. 5, which consists of two coupled neural networks [34–36]. The first feedforward neural network ***µ*** models the nervous system of the fish (actor). The actor network ***µ*** implements the control and aims to estimate the optimal control **u**^∗^. The second feedforward neural network *Q* models the hunger cues of the fish (critic). The critic network *Q* determines whether the implemented control ***µ*** is optimal and aims to estimate the negative interception radius *−R*.

**FIG 5:**
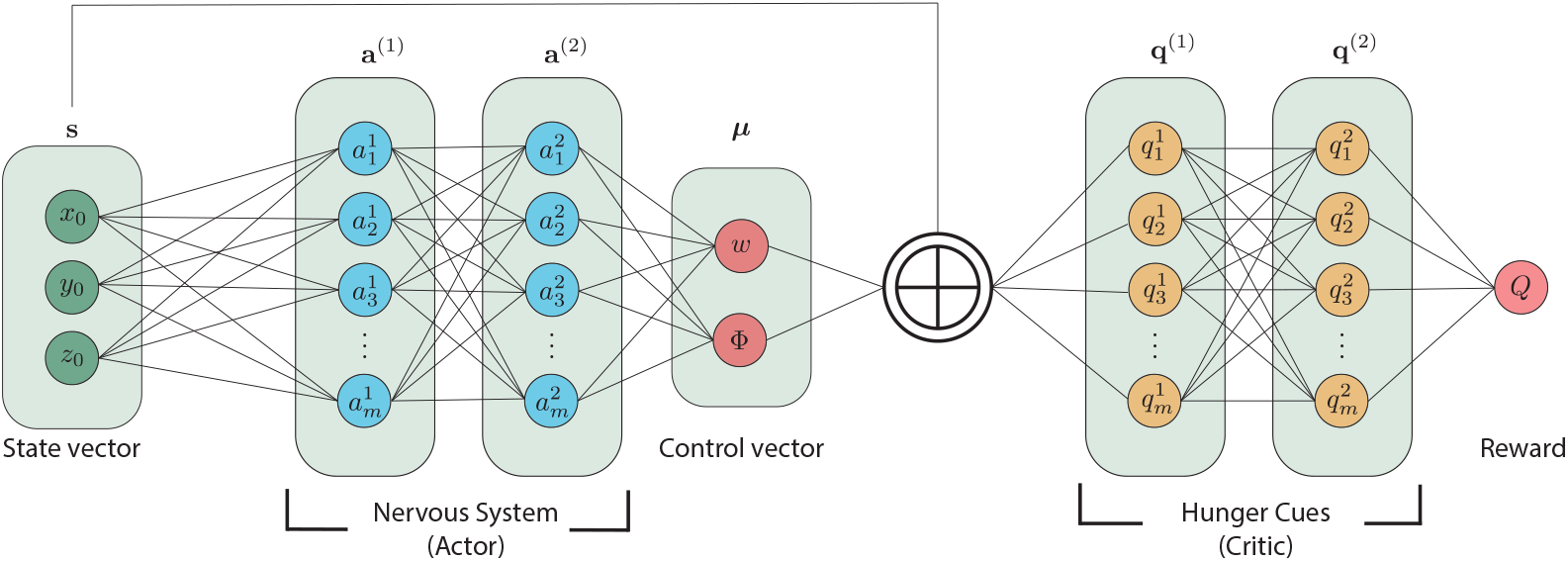
Schematic of the deep neural network used for reinforcement learning with the actor-critic paradigm. The actor network models the nervous system of the fish which, given an initial state **s**, outputs a control ***µ*** to be actuated. The initial state and the actor network output is then inputted to the critic network, modeling the hunger cues in the fish, which outputs the reward *Q* for the interception. The action to be learned by the actor network is **u**^∗^ and the reward to be learned by the critic network is the negative interception radius *−R*.

During repeated one-shot interceptions, the fish learns to implement the optimal control policy by appraising the result of an interception using a value function. An optimal one-shot interception has a noiseless optimal control policy given by **u**^∗^ = (1 *− ρ*_0_*/*(*vT*), *ϕ*_0_) from Eqs. (12), (14), and value function given by the magnitude of the interception error in Eq. (7)^4^.

The actor network ***µ*** has input layer given by the initial state **s**, two hidden layers (64 neurons each, ReLU activation), and output layer obtained by bifurcating the final hidden layer into two separate outputs, one for the wait fraction *w* (sigmoid activation) and the other for the heading angle Φ (hyperbolic tangent activation). The two outputs are concatenated to form the actor network output,

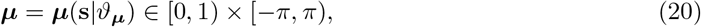

Where *v*_***µ***_ are the actor network parameters.

The critic network has input layer given by a state-control pair **s** *⊕* ***µ*** *∈* ℝ3 *×* [0, 1) *×* [ *−π, π*), two hidden layers (64 neurons each, ReLU activation) and output layer *Q* (identity activation function) given by

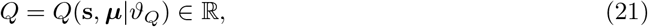

where *θ*_*Q*_ are the parameters the critic network. The critic network output *Q* is dense and rewards near misses to promote parameter learning at early times when interception accuracy is low.

## VII. DEEP REINFORCEMENT LEARNING RESULTS

Initially, random parameters are chosen for *v*_***µ***_, *v*_*Q*_. The interceptions are conducted in a 10 *×* 10 box in which the distance separating any two points selected at random is approximately 5.2 [38]. The interception error at episode 1 is approximately 5.5, which is larger than 5.2 due to a finite sample size. There is an initial trial-and-error period where the fish is attempting interceptions that are not successful with an error of approximately 5.5 *−* 6.5, larger than the error of 5.5 given by random weights (Fig. 6a). Then, after several hundred episodes, the interception error begins decreasing at a near constant rate (Fig. 6b). At this stage, the network is learning the optimal parameters for the interception. After around 2500 episodes, the interception error is approximately 1, which we define as the network having learned the optimal polices for feeding (Fig. 6c).

**FIG 6:**
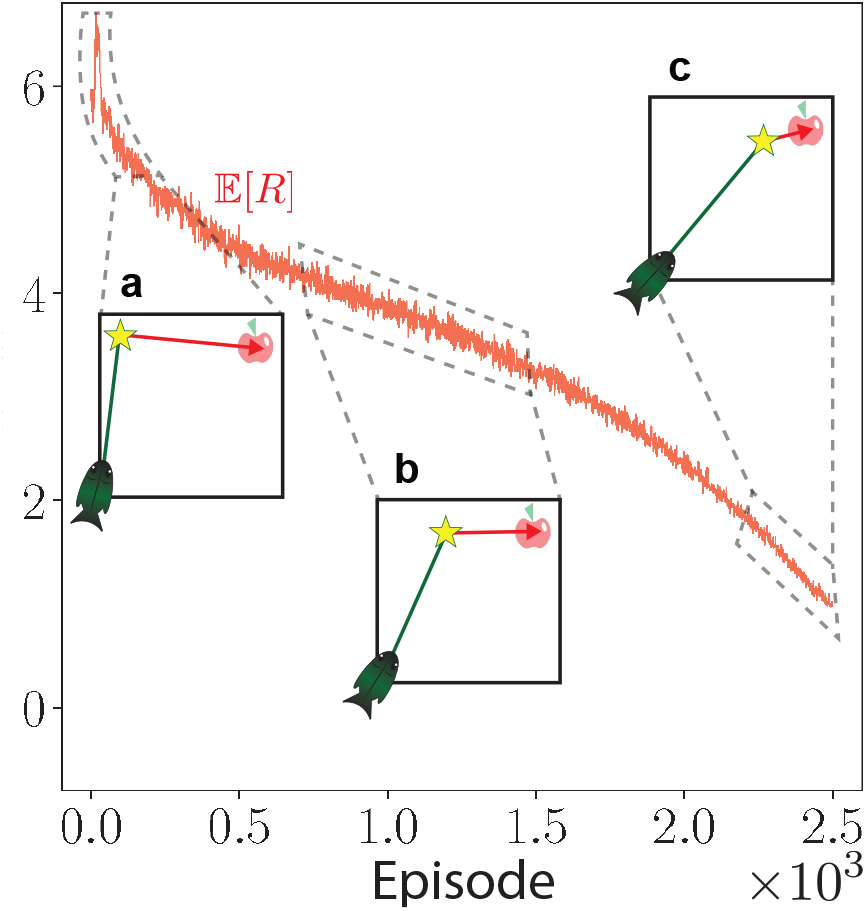
Deep reinforcement learning error during episodic training. The average interception error (red curve) is shown over the batch of 1000 interception replicates for each episode. Insets show schematic illustrations of different stages of learning. (a) Initial trial-and-error period; (b) improvement; (c) final trained agent.

The learning of the negative interception radius *−R* and the optimal control **u**^∗^ are coupled during episodic training as the neural networks for actor and critic are coupled, which couples updating the parameters *v*_***µ***_, *v*_*Q*_. The fish learns how to appraise a given control before learning how to implement controls which provide the highest rewards. The actor network output ***µ*** = (*w*_*RL*_, Φ_*RL*_) during episodic training is compared to the optimal control **u**^∗^ in Fig. 7. Initially, at episode 1, the correlation between the actor network output and the optimal control is very low. By episode 1000, the heading angle Φ_*RL*_ is well correlated with the optimal heading angle *ϕ*_0_, showing that the actor network first learns to aim in the direction of the impact point of the fruit. By episode 2000, the wait fraction *w*_*RL*_ is well correlated with the optimal wait fraction 1 *− ρ*_0_*/*(*vT*).

**FIG 7:**
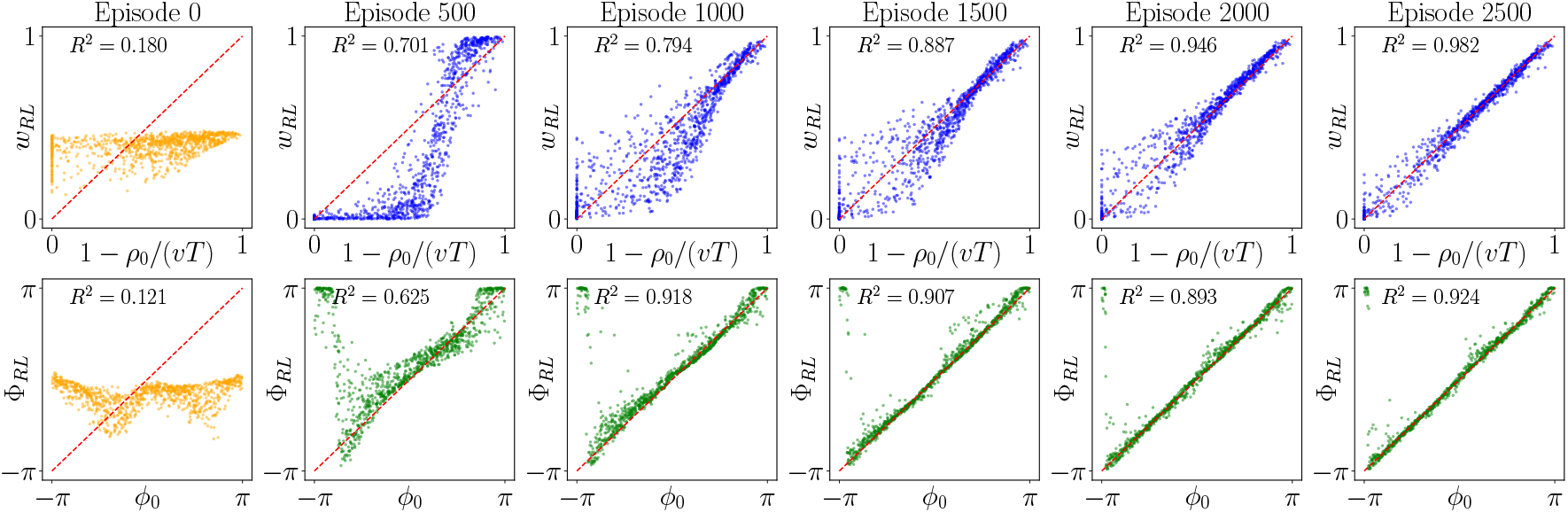
Actor network output ***µ*** = (*w*_*RL*_, Φ_*RL*_) and optimal control **u**^∗^ = (1 *−* (*ρ*_0_*/vT*), *ϕ*_0_) during episodic reinforcement learning. There are 1000 replicates for each episode shown. Initially, the network parameters were chosen randomly (orange points) and training occurred for 2500 episodes. *R*^2^ is the coefficient of determination for the wait fraction and the circular correlation coefficient [37] for the heading angle.

## VIII. DISCUSSION AND OUTLOOK

Inspired by fruit-catching behavior in *B. guatemalensis*, we have presented a model for the optimal control and learning in one-shot interceptions where the timescale of target dynamics during the interception is on the order of the time for the agent to process and act on sensory information. Our analytic results show a non-vanishing optimal interception error in the stochastic and kinetically-penalized regime which is consistent with experimental observations.

RL simulations complement our analytic approaches and show that learning occurs stepwise: first the form of the error of an interception is learned, after which the agent learns to aim in the direction of the impact point of the target and finally the agent learns the optimal wait time to intercept the target.

The basic features of one-shot interceptions are probably seen at the end of any adaptive interception that involves continuous feedback. This occurs because during the final update of the control process, the timescales of sensing and action are so close as to be an open-loop problem similar to that in a one-shot interception. Future work would aim to generalize the model presented in this paper for more realistic situations, including those with dependent sources of noise sharing a joint probability distribution and incorporating air resistance into the model. We also anticipate future experimental work may investigate the role of cooperation and competition for one-shot interceptions with multiple agents, such as in a school of fish. Generalizing our ideas to both natural and artificial settings involving landing, perching and other critical transitions might be a natural next step.

## ACKNOWLEDGMENTS

This research was inspired by early work conducted by Andrew Marantan as part of his PhD thesis [39]. The authors gratefully acknowledge helpful discussions with Andrew Marantan during the conceptualization and early development of this study, and thank Stefan Schuster for helpful discussions and feedback on experimental *B. guatemalensis* interceptions.

## FUNDING

Aria Azari-Pour is supported by the Herchel Smith Fund of the University of Cambridge through a Herchel Smith Fellowship and an Exchange Scholar Award at Harvard University.

## ADDITIONAL INFORMATION

### Data and code availability

The code used to generate the figures in this paper is available online [40].

### Author contributions

A. A.: conceptualization, investigation, methodology, formal analysis, software validation, funding acquisition, writing - original draft, writing - reviewing and editing; L. M.: conceptualization, supervision, validation, writing - reviewing and editing.

### Conflict of interest statement

The authors declare no competing interests.

## Footnotes

1 The cost function in Eq. (4) does not contain a running cost as u is constant during the interception. One may add a term proportional to ∥u∥ to penalize singularities in the control; however, since ∥u∥ is bounded as w is bounded and Φ is periodic, we do not consider such a term.

2 One may also consider noise in *T*, which is equivalent to noise in the measurement of *z*0 or g by the fish. In the following, we do not consider noise in *T* for simplicity.

3 The inertial term has the form of the time-integral of the kinetic energy of the fish over the fraction of time that the fish is swimming.

4 An interception may be represented by a Markov decision process, which lends well to the use of RL methods for learning optimal policies and rewards [34].

## Notes

https://github.com/aa2479/FruitFishRL

### Competing Interest Statement

The authors have declared no competing interest.

## References

[1] M. Mischiati, H.-T. Lin, P. Herold, E. Imler, R. Olberg, and A. Leonardo, Nature 517, 333 (2015).

[2] P. Gerullis and S. Schuster, Current Biology 24, 2156 (2014).

[3] K. Ghose, T. K. Horiuchi, P. S. Krishnaprasad, and C. F. Moss, PLoS biology 4, e108 (2006).

[4] S. A. Kane, A. H. Fulton, and L. J. Rosenthal, The Journal of Experimental Biology 218, 212 (2015).

[5] N. Hogan, IEEE Transactions on Automatic Control 29, 681 (1984).

[6] N. Hogan, The Journal of Neuroscience 4, 2745 (1984).

[7] M. K. McBeath, D. M. Shaffer, and M. K. Kaiser, Science 268, 569 (1995).

[8] F. Lacquaniti, G. Bosco, I. Indovina, B. La Scaleia, V. Maffei, A. Moscatelli, and M. Zago, Frontiers in Integrative Neuroscience 7, 10.3389/fnint.2013.00101 (2013).

[9] T. Haarnoja, B. Moran, G. Lever, S. H. Huang, D. Tirumala, J. Humplik, M. Wulfmeier, S. Tunyasuvunakool, N. Y. Siegel, R. Hafner, M. Bloesch, K. Hartikainen, A. Byravan, L. Hasenclever, Y. Tassa, F. Sadeghi, N. Batchelor, F. Casarini, S. Saliceti, C. Game, N. Sreendra, K. Patel, M. Gwira, A. Huber, N. Hurley, F. Nori, R. Hadsell, and N. Heess, Science Robotics 9, 10.1126/scirobotics.adi8022 (2024).

[10] D. A. Haggerty, M. J. Banks, E. Kamenar, A. B. Cao, P. C. Curtis, I. Mezić, and E. W. Hawkes, Science Robotics 8, 10.1126/scirobotics.add6864 (2023).

[11] P. Cigliano, V. Lippiello, F. Ruggiero, and B. Siciliano, IEEE Transactions on Control Systems Technology 23, 1657 (2015).

[12] P. Androulakakis, Z. E. Fuchs, and J. E. Shroyer, in 2018 IEEE Congress on Evolutionary Computation (CEC) (IEEE, 2018) pp. 1–9.

[13] P. Dürr, M. El Gheche, G. J. Maeda, N. Mukai, N. Takahashi, S. Heusser, H. Sahloul, Y. Saraiji, P. Adodin, Y. Bi, S. Blakeman, C. Conti, D. Fuentes Hitos, Y. Hu, F. Khadivar, R. Kreiser, L. Martinez, F. Schilling, R. Tapiador Morales, G. Torrente, M. Ynocente Castro, L. Abecassis, A. Giammarino, Y. T. Huang, Y. Nagel, A. Scotti, A. Sigrist, T. Silva, E. Walther, J. Wong, B. Yang, A. Aydin, D. Grover, A. Saha, V. Cavinato, T. Kakinuma, T. Kunori, V. Monferrato, S. Richter, S. Charalambous, S. Guist, M. A. Kuhlmann-Jorgensen, L. Miele, A. Politis, M. Scardecchia, H. Kitano, P. R. Wurman, P. Stone, and M. Spranger, Nature 652, 886 (2026).

[14] V. Shaferman and T. Shima, Journal of Guidance, Control, and Dynamics 38, 1395 (2015).

[15] V. Fomin, Optimal Filtering (Springer Netherlands, Dordrecht, 1999).

[16] Q. Wang, C. Ye, G. Chang, C. Tang, and X. Zhang, Scientific Reports 10.1038/s41598-026-50388-3 (2026).

[17] A. C. Noel, H.-Y. Guo, M. Mandica, and D. L. Hu, Journal of The Royal Society Interface 14, 20160764 (2017).

[18] R. Jacquot, K. Ben Mansour, K. Bouillet, J.-P. Jehl, and G. Gauchard, Frontiers in Sports and Active Living 7, 10.3389/fspor.2025.1650300 (2025).

[19] K. E. Drewe, M. H. Horn, K. A. Dickson, and A. Gawlicka, Journal of Fish Biology 64, 890 (2004).

[20] M. H. Horn, Oecologia 109, 259 (1997).

[21] P. Krupczynski and S. Schuster, Current Biology 18, 1961 (2008).

[22] J. Gray, Proceedings of the Royal Society of London. Series B, Containing Papers of a Biological Character 113, 115 (1933).

[23] W. C. Witt, L. Wen, and G. V. Lauder, Integrative and Comparative Biology 55, 728 (2015).

[24] S. Schuster, Journal of Comparative Physiology A 209, 827 (2023).

[25] R. S. Sutton and A. G. Barto, Reinforcement learning : an introduction (MIT Press, 1998).

[26] G. B. Margolis, G. Yang, K. Paigwar, T. Chen, and P. Agrawal, The International Journal of Robotics Research 43, 572 (2024).

[27] R. Chen and J. H. Goldberg, Current Opinion in Neurobiology 65, 1 (2020).

[28] J. Kasdin, A. Duffy, N. Nadler, A. Raha, A. L. Fairhall, K. L. Stachenfeld, and V. Gadagkar, Nature 641, 699 (2025).

[29] E. D. Tytell and G. V. Lauder, Journal of Experimental Biology 205, 2591 (2002).

[30] M. Gazzola, W. M. Van Rees, and P. Koumoutsakos, Journal of Fluid Mechanics 698, 5 (2012).

[31] C. M. Bishop and H. Bishop, Deep Learning (Springer International Publishing, Cham, 2024).

[32] B. Sun, S. Mohamed Haris, and R. Ramli, Instruments 10, 8 (2026).

[33] C. Tang, B. Abbatematteo, J. Hu, R. Chandra, R. Martín-Martín, and P. Stone, Annual Review of Control, Robotics, and Autonomous Systems 8, 153 (2025).

[34] I. Grondman, L. Busoniu, G. A. D. Lopes, and R. Babuska, IEEE Transactions on Systems, Man, and Cybernetics, Part C (Applications and Reviews) 42, 1291 (2012).

[35] K. Arulkumaran, M. P. Deisenroth, M. Brundage, and A. A. Bharath, IEEE Signal Processing Magazine 34, 26 (2017).

[36] I. Drori, The Science of Deep Learning (Cambridge University Press, 2022).

[37] S. R. Jammalamadaka and Y. R. Sarma, in Statistical Theory and Data Analysis II: Proceedings of the Second Pacific Area Statistical Conference, edited by K. Matusita (North-Holland, Amsterdam, 1988) pp. 349–364.

[38] B. Ghosh, Bulletin of Calcutta Mathematical Society 43, 17 (1951).

[39] A. W. Marantan, The Roles of Randomness in Biophysics: From Cell Growth to Behavioral Control, Ph.D. thesis, Harvard University (2017).

[40] A. Azari-Pour, Modeling one-shot interceptions in fruit-catching fish (2026).

