## Supplementary Information for "Modeling one-shot interceptions in fruit-catching fish"

#### I. DERIVATION OF STOCHASTIC COST FUNCTION

In this section we derive the form of the cost function used for the fruit-fish interception. The inertial term is chosen as the integral of the kinetic energy of the fish with unit mass over the swim fraction,

$$\mathcal{Q}(v; \mathbf{s}, \mathbf{u}) = \int_{wT}^T \frac{1}{2} v^2 dt = \frac{1}{2} v^2 (1 - w) T. \quad (1)$$

The interception error, with noise in the state $\mathbf{s}$ and the control $\mathbf{u}$, is

$$\mathbf{e}(\mathbf{s}, v, \mathbf{u}) = \begin{bmatrix} x_0 + \xi_1 - v(1 - w)T \cos(\Phi + \eta) \\ y_0 + \xi_2 - v(1 - w)T \sin(\Phi + \eta) \\ 0 \end{bmatrix}. \quad (2)$$

The squared stochastic error is

$$\mathbf{e}^T \mathbf{e} = [x_0 + \xi_1 - v(1 - w)T \cos(\Phi + \eta)]^2 + [y_0 + \xi_2 - v(1 - w)T \sin(\Phi + \eta)]^2, \quad (3)$$

$$= \rho_0^2 + v^2(1 - w)T^2 + \|\boldsymbol{\xi}\|^2 - 2v(1 - w)T \left[ x_0 \cos(\Phi + \eta) + y_0 \sin(\Phi + \eta) \right] + \mathcal{F}(\boldsymbol{\xi}, \eta), \quad (4)$$

where $\|\boldsymbol{\xi}\|^2 = \xi_1^2 + \xi_2^2$ and we define

$$\mathcal{F}(\boldsymbol{\xi}, \eta) = 2(x_0 \xi_1 + y_0 \xi_2) - 2v(1 - w)T \left[ \xi_1 \cos(\Phi + \eta) + \xi_2 \sin(\Phi + \eta) \right], \quad (5)$$

which is a stochastic function coupling $\boldsymbol{\xi}$ and $\eta$. Using $(x_0, y_0) = \rho_0(\cos \phi_0, \sin \phi_0)$, we may write

$$\mathbf{e}^T \mathbf{e} = \rho_0^2 + v^2(1 - w)T^2 + \|\boldsymbol{\xi}\|^2 + \mathcal{F}(\boldsymbol{\xi}, \eta) - 2v(1 - w)T \rho_0 \left[ \cos(\Phi + \eta) \cos \phi_0 + \sin(\Phi + \eta) \sin \phi_0 \right], \quad (6)$$

$$= \rho_0^2 + v^2(1 - w)T^2 + \|\boldsymbol{\xi}\|^2 + \mathcal{F}(\boldsymbol{\xi}, \eta) - 2v(1 - w)T \rho_0 \left[ \cos \eta \cos(\Phi - \phi_0) - \sin \eta \sin(\Phi - \phi_0) \right], \quad (7)$$

where we have used the identity in Eq. (7) for any two angles $\varphi_1, \varphi_2 \in [-\pi, \pi)$,

$$\cos(\varphi_1 - \varphi_2) = \cos \varphi_1 \cos \varphi_2 + \sin \varphi_1 \sin \varphi_2. \quad (8)$$

Assume $\boldsymbol{\xi}$ and $\eta$ satisfy

$$\mathbb{E}[f(\boldsymbol{\xi})g(\eta)] = \mathbb{E}[f(\boldsymbol{\xi})]\mathbb{E}[g(\eta)], \quad (9)$$

for any smooth functions $f, g$ where $\mathbb{E}[f(\boldsymbol{\xi})], \mathbb{E}[g(\eta)] < \infty$. As $\mathbb{E}[\boldsymbol{\xi}] = 0$, and $\mathcal{F}$ is a linear combination of $\xi_1, \xi_2$, then we must have

$$\mathbb{E}[\mathcal{F}(\boldsymbol{\xi}, \eta)] = 0. \quad (10)$$

\*

†

The probability distribution of $\eta$, $p(\eta)$, is symmetric about $\eta = 0$ as the error in the heading angle of the fish has equal probability to be in the clockwise- or counterclockwise-direction. It follows that

$$\mathbb{E}[\sin \eta] = \int_{-\pi}^{\pi} \sin \eta p(\eta) d\eta = 0, \quad (11)$$

as the integrand is odd and the domain of integration is symmetric about $\eta = 0$.

The cost function in the fruit-fish interception then follows from Eqs. (10) and (11), as well as $\sigma_{\xi}^2 = \mathbb{E}[\|\xi\|^2]$.

TABLE I: List of terms used in the main paper.

| Term | Description |
| --- | --- |
| $\mathbf{x}(t)$ | Trajectory of target with components $x_i(t)$ for $i = 1, 2, 3$ |
| $\mathbf{y}(t; \mathbf{u})$ | Trajectory of agent with components $y_i(t)$ for $i = 1, 2, 3$ (depends on $v, \mathbf{u}$) |
| $T$ | Impact time for the target to reach the plane $z = 0$ |
| $g$ | Acceleration due to gravity |
| $w \in [0, 1)$ | Wait fraction control input actuated by the agent |
| $\Phi \in [-\pi, \pi)$ | Heading angle control input actuated by the agent |
| $\mathbf{u} = (w, \Phi)$ | Control actuated by agent (constant over $[0, T]$) |
| $v$ | Launch velocity of agent (constant over $[0, T]$) |
| $\mathbf{u}^* = (w^*, \Phi^*)$ | Optimal control actuated by agent |
| $\mathbf{r}(t; \mathbf{u})$ | Relative position of the target with respect to the agent given by $\mathbf{x}(t) - \mathbf{y}(t; \mathbf{u})$ |
| $\mathbf{s} = \mathbf{r}(0)$ | Initial state of the interception at time $t = 0$ |
| $\mathbf{e} = \mathbf{r}(T; \mathbf{u})$ | Terminal error of the interception at time $t = T$ |
| $\mathcal{J}$ | Stochastic cost function to be minimized to find $\mathbf{u}^*$ (depends on $\mathbf{e}, \mathbf{s}, v, \mathbf{u}$) |
| $\mathcal{Q}$ | Inertial penalty in the cost function $\mathcal{J}$ |
| $\beta$ | Hunger parameter (encourages interception) |
| $M$ | Inertia (fatigue) parameter |
| $\alpha = M/\beta$ | Laziness parameter of the agent |
| $\mathcal{W}(t)$ | Cumulative time the fruit-fish has been swimming at time $t$ |
| $(x_0, y_0, z_0)$ | Initial state $\mathbf{s}$ of the fruit-fish interception |
| $\rho_0$ | Distance from the fruit-fish to the impact point, equivalent to $(x_0^2 + y_0^2)^{1/2}$ |
| $\rho^*$ | Optimal distance traveled by the fish, equivalent to $(1 - w^*)vT$ |
| $\xi = (\xi_1, \xi_2, 0)$ | Additive noise in the measurement of the impact point $(x_0, y_0, 0)$ by the fish |
| $\eta$ | Additive noise in the actuation of the heading angle control input $\Phi$ by the fish |
| $\sigma_X^2$ | Variance of the random variable $X$ |
| $C_\eta = \mathbb{E}[\cos \eta]$ | Attenuation factor for fruit-fish interceptions from heading angle noise |
| $k$ | Variance decay rate of $\sigma_\xi$ as a function of $w$ at $w^*$ |
| $R$ | Interception radius, equivalent to $\Vert \mathbf{e}\Vert $ |
| $\varepsilon$ | Tolerance of the fish for lateral deflections due to heading angle noise |
| $\mathcal{P}(\varepsilon)$ | Probability that the fish misses the fruit due to heading angle noise |
| $p(X)$ | Probability distribution function of a random variable $X$ |
| $F(X)$ | Cumulative distribution function of a random variable $X$ |
| $\vartheta_\mu$ | Learning parameters (weights and biases) of the actor network $\mu$ |
| $\vartheta_Q$ | Learning parameters (weights and biases) of the critic network $Q$ |
| $\mu = (w_{RL}, \Phi_{RL})$ | Actor network output, an estimator for $\mathbf{u}^*$ in the noiseless limit |
| $Q$ | Critic network output, an estimator for $-R$ |

### II. NEURAL NETWORK ARCHITECTURE

The deep learning architecture used in this work is described in this section. Each layer $i = 0, 1, 2, \dots, N + 1$ consists of $m_i$ neurons and each neuron $j \in m_i$ has activation $a_j^{(i)} \in \mathbb{R}$. We denote layer $i = 0$ as the input layer, layer $i = N + 1$ as the output layer, and the $N$ layers $i = 1, \dots, N$ as the hidden layers. The activation of neuron $j$ in layer $i + 1$ is given by a linear combination of the $m_i$ neuron activations in the previous layer,

$$a_j^{(i+1)} = \mathcal{A}_{i+1} \left( \sum_{k=1}^{m_i} w_{jk}^{(i+1)} a_k^{(i)} + b_j^{(i+1)} \right), \quad (12)$$

with $\mathcal{A}_{i+1}$ a potentially non-linear activation function for layer $i + 1$. In Eq. (12), $w_{jk}^{(i+1)}$ is the synaptic weight connecting neuron $k$ in layer $i$ to neuron $j$ in layer $i + 1$, and $b_j^{(i+1)}$ is the bias of neuron $j$ in layer $i + 1$. Let $\mathbf{W}^{(i+1)} = [\mathbf{w}_1^{(i+1)}, \dots, \mathbf{w}_{m_i}^{(i+1)}] \in \mathbb{R}^{m_{i+1} \times m_i}$ be the weight matrix from layer $i$ to layer $i + 1$ with $\mathbf{w}_j^{(i+1)}$ the synaptic weights from the $m_i$ neurons in layer $i$ to neuron $j$ in layer $i + 1$. Then, if $\mathbf{a}^{(i)} = (a_1^{(i)}, \dots, a_{m_i}^{(i)})$ is the vector of neuron activations in layer $i$ and $\mathbf{b}^{(i)} = (b_1^{(i)}, \dots, b_{m_i}^{(i)})$ are the biases, the activation of layer $i + 1$ given layer $i$ is

$$\mathbf{a}^{(i+1)} = \mathcal{A}_{i+1}(\mathcal{R}_{i+1}[\mathbf{a}^{(i)}]), \quad (13)$$

where

$$\mathcal{R}_{i+1}[\mathbf{a}^{(i)}] = \mathbf{W}^{(i+1)} \mathbf{a}^{(i)} + \mathbf{b}^{(i)}, \quad (14)$$

is an affine transformation from layer $i$ to layer $i + 1$, and $\mathcal{A}_{i+1}$ is the corresponding activation function. Letting $\mathcal{T}_{i+1} = \mathcal{A}_{i+1} \circ \mathcal{R}_{i+1}$, then the output $\mathbf{a}^{(N+1)}$ is

$$\mathbf{a}^{(N+1)} = (\mathcal{T}_{N+1} \circ \dots \circ \mathcal{T}_2 \circ \mathcal{T}_1)(\mathbf{a}^{(0)}). \quad (15)$$

For the network $\mathcal{T}_{N+1} \circ \dots \circ \mathcal{T}_2 \circ \mathcal{T}_1$, let

$$\vartheta = \bigoplus_{(i,j,k) \in \mathcal{I}} (w_{jk}^{(i+1)}, b_j^{(i+1)}), \quad (16)$$

be the vector of weights and biases in the network, which we refer to as the *parameters* of the network, with index set $\mathcal{I} \subset \mathbb{Z}^3$, which has dimension

$$\dim \vartheta = \prod_{i=0}^N m_{i+1} (m_i + 1), \quad (17)$$

or, if the number of neurons in each of the hidden layers is $m$,

$$\dim \vartheta = m(m_0 + 1) + m_{N+1}(m + 1) + [m(m + 1)]^{N-1}. \quad (18)$$

### III. ACTOR NETWORK ARCHITECTURE

The actor network $\boldsymbol{\mu}$ has input layer $\mathbf{s}$ and output layer $\boldsymbol{\mu} \in [0, 1) \times [-\pi, \pi)$. There are two hidden layers, each with $m = 64$ neurons. Letting $\vartheta_{\boldsymbol{\mu}}$ denote the parameters of the actor network, the network is formally,

$$\boldsymbol{\mu} = \boldsymbol{\mu}(\mathbf{s} | \vartheta_{\boldsymbol{\mu}}) = (\mathcal{L}_3 \circ \mathcal{L}_2 \circ \mathcal{L}_1)(\mathbf{s}), \quad (19)$$

where the layer transformations $\mathcal{L}_1, \mathcal{L}_2$ are

$$\mathbf{a}^{(i+1)} = \mathcal{L}_{i+1}(\mathbf{a}^{(i)}) = \text{ReLU}(\mathbf{W}^{(i+1)} \mathbf{a}^{(i)} + \mathbf{b}^{(i+1)}), \quad (20)$$

with weights $\mathbf{W}^{(1)} \in \mathbb{R}^{m \times 3}$, $\mathbf{W}^{(2)} \in \mathbb{R}^{m \times m}$, and biases $\mathbf{b}^{(1)}, \mathbf{b}^{(2)} \in \mathbb{R}^m$, where $\text{ReLU}(\cdot)$ is the rectilinear unit.

For the final layer transformation $\mathcal{L}_3$ which outputs the control $\boldsymbol{\mu}$, we split the transformation into two scalar transformations for the wait fraction $w$ and heading angle $\Phi$. The final layer transformation may be written as

$$\boldsymbol{\mu} = \mathcal{L}_3(\mathbf{a}^{(2)}) = \sigma(\mathcal{R}_{3,w} \mathbf{a}^{(2)}) \oplus \tanh(\mathcal{R}_{3,\Phi} \mathbf{a}^{(2)}), \quad (21)$$

where $\sigma(\cdot)$ is the sigmoid function and $\tanh(\cdot)$ is the hyperbolic tangent function. The affine transformations for the final layer are $\mathcal{R}_{3,\alpha} \mathbf{a}^{(2)} = \mathbf{W}^{(3,\alpha)} \mathbf{a}^{(2)} + \mathbf{b}^{(3,\alpha)}$ with weight matrices $\mathbf{W}^{(3,\alpha)} \in \mathbb{R}^{1 \times m}$ and bias $\mathbf{b}^{(3,\alpha)} \in \mathbb{R}$ for each of the controls $\alpha = w, \Phi$.

##### IV. CRITIC NETWORK ARCHITECTURE

The critic network $Q$ has input layer $\mathbf{s} \oplus \boldsymbol{\mu}$, where $\mathbf{s}$ is the input layer of the actor network and $\boldsymbol{\mu} = \boldsymbol{\mu}(\mathbf{s}|\vartheta_{\boldsymbol{\mu}})$ is the output layer of the actor network. There are two hidden layers, each consisting of $m = 64$ neurons, and the output layer is the scalar reward $Q \in \mathbb{R}$. The critic network is formally,

$$Q = Q(\mathbf{s}, \boldsymbol{\mu}|\vartheta_Q) = (\mathcal{H}_3 \circ \mathcal{H}_2 \circ \mathcal{H}_1)(\mathbf{s} \oplus \boldsymbol{\mu}), \quad (22)$$

with $\vartheta_Q$ the learning parameters of the critic network. The layer transformations $\mathcal{H}_i$ have the same form as Eq. (20) with ReLU activation functions for the two hidden layers and the identity activation function for the final output layer. The weight matrices are $\mathbf{W}^{(1)} \in \mathbb{R}^{m \times 5}$, $\mathbf{W}^{(2)} \in \mathbb{R}^{m \times m}$, $\mathbf{W}^{(3)} \in \mathbb{R}^{1 \times m}$ and the biases are $\mathbf{b}^{(1)} \in \mathbb{R}^m$, $\mathbf{b}^{(2)} \in \mathbb{R}^m$, $\mathbf{b}^{(3)} \in \mathbb{R}$.

##### V. SIMULATION ENVIRONMENT

The motion of the fish is confined onto the spatial domain $[0, L]^2 = [0, L] \times [0, L] \subset \mathbb{R}^2$ where we choose $L = 10$ in the simulations. The initial position of the fish $\mathbf{y}(0)$ and the impact point of fruit $\mathbf{x}(T)$ are sampled uniformly and independently over $[0, L]^2$ as $\mathbf{x}(T), \mathbf{y}(0) \sim \mathcal{U}([0, L]^2)$, where $\mathcal{U}(\Omega)$ is the uniform distribution over the set $\Omega \subset \mathbb{R}^d$ in $d$ dimensions. The impact time is sampled from a uniform distribution $T \sim \mathcal{U}([1.0, 6.0])$. In the simulations, the fish speed is fixed as $v = 3.5$, and therefore a physical interception requires $3.5T \geq \rho_0$, where $\rho_0 = \|\mathbf{x}(T) - \mathbf{y}(0)\|$. For some values of $T$ and $\rho_0$, the inequality is not satisfied and the interception is not possible. In this case, we retain this simulation to train the neural network. We chose the wait time as $\max(0, w)$ such that if the interception is impossible, with $w < 0$, the fish swims immediately and learns the heading angle. At the beginning of the simulation, the input of the actor network is $\mathbf{s} = \mathbf{x}(T) - \mathbf{y}(0)$. At the end of the simulation when $t = T$, we constrain the position of the fish to be inside the box $[0, L]^2$.

##### VI. NEURAL NETWORK TRAINING

For each of the $\tau = 1, \dots, 2500$ episodes, a full forward execution of the episode consists of three operations occurring sequentially. First, there are $N = 1000$ independent interception scenarios generated as a batch. For each scenario $k = 1, \dots, N$, a random state $\mathbf{s}_k^{(\tau)}$ is constructed. Second, the $N$ states are inputted as a batch into the actor network with parameters $\vartheta_{\boldsymbol{\mu}}$ and the network computes the estimate of the control with additive Gaussian noise,

$$\boldsymbol{\mu}_k^{(\tau)} = \boldsymbol{\mu}(\mathbf{s}_k^{(\tau)}|\vartheta_{\boldsymbol{\mu}}^{(\tau)}) + \mathcal{N}(0, \Sigma(\tau)), \quad (23)$$

where $\Sigma(\tau)$ is the *exploration variance* which is a monotone decreasing function of $\tau$. At early episodes, the noise encourages broad exploration to prevent the network from being trapped in local minima, whereas at later episodes the network shifts to using the learned optimal policy. Third, the stochastic actor network output $\boldsymbol{\mu}_k^{(\tau)}$ is used to calculate the resulting trajectories, the interception error magnitude $R_k^{(\tau)}$, and the critic network output $Q_k^{(\tau)}$ with the critic network parameters $\vartheta_Q^{(\tau)}$ for each of the $N$ fish.

Following the full forward execution of each episode, the network parameters $\vartheta_{\boldsymbol{\mu}}, \vartheta_Q$ are updated. For the critic network, we define the critic loss function as the mean-squared error of the critic network output layer over the batch of size $N$,

$$\mathcal{L}_Q(\vartheta_Q^{(\tau)}) = \frac{1}{N} \sum_{k=1}^N \left[ Q(\mathbf{s}_k^{(\tau)}, \boldsymbol{\mu}_k^{(\tau)}|\vartheta_Q^{(\tau)}) + R_k^{(\tau)} \right]^2. \quad (24)$$

The critic network parameters are updated to minimize the critic loss in Eq. (24) as

$$\vartheta_Q^{(\tau+1)} \leftarrow \underset{\vartheta_Q^{(\tau)}}{\operatorname{argmin}} \mathcal{L}_Q(\vartheta_Q^{(\tau)}). \quad (25)$$

Minimization of Eq. (24) occurs by computing the gradient of the critic loss through backpropagation and then updating $\vartheta_Q$ in the direction opposite to the gradient of the critic loss using the Adam optimization algorithm.

We define the actor network loss function as the negative mean of the critic network output for fixed $\vartheta_Q$,

$$\mathcal{L}_\mu(\vartheta_\mu^{(\tau)}) = -\frac{1}{N} \sum_{k=1}^N Q(\mathbf{s}_k^{(\tau)}, \mu(\mathbf{s}_k^{(\tau)} | \vartheta_\mu^{(\tau)}) | \vartheta_Q^{(\tau+1)}). \quad (26)$$

The actor network parameters are updated to minimize the actor loss in Eq. (26) as

$$\vartheta_\mu^{(\tau+1)} \leftarrow \underset{\vartheta_\mu^{(\tau)}}{\operatorname{argmin}} \mathcal{L}_\mu(\vartheta_\mu^{(\tau)}). \quad (27)$$

Minimization of Eq. (26) occurs through backpropagation and the chain rule through the critic network. The parameters $\vartheta_\mu$ are updated such that the actions produced from the updated parameters $\vartheta_\mu$ result in the maximum possible estimated reward $Q$ from the critic network.

**Algorithm 1** Deep Reinforcement Learning Episodic Training

```
1: Input Total number of episodes, $E$
2: Input Batch size per episode, $N$
3: Input Initial and final exploration noise standard deviations, $\sigma_1, \sigma_E$
4: Input Learning rates for actor and critic networks, $\eta_\mu, \eta_Q$

5: Initialize Actor network parameters $\vartheta_\mu$ with random entries
6: Initialize Critic network parameters $\vartheta_Q$ with random entries
7: Initialize Adam optimizer for actor with learning rate $\eta_\mu$
8: Initialize Adam optimizer for critic with learning rate $\eta_Q$

9: for episode $\tau = 1$ to $E$ do
10:   // 1. Environment and state generation
11:   $\sigma \leftarrow \sigma_1 - (\sigma_1 - \sigma_E)(\tau - 1)/(E - 1)$ ▷ Linearly anneal exploration noise
12:   Generate batch of $N$ impact points $\mathbf{x}(T)$ uniformly on $[0, 10]^2 \subset \mathbb{R}^2$
13:   Generate batch of $N$ initial fish positions $\mathbf{y}(0)$ uniformly on $[0, 10]^2 \subset \mathbb{R}^2$
14:   Generate batch of $N$ impact times $T$ uniformly on $[1, 6] \subset \mathbb{R}$
15:   $\mathbf{s} \leftarrow (\mathbf{x}(T) - \mathbf{y}(0)) \oplus T$ ▷ Use impact time $T$ instead of $z$ for simplicity

16:   // 2. Action selection with exploration
17:   $\mu_{pred} \leftarrow \mu(\mathbf{s} | \vartheta_\mu)$ ▷ Forward execution through actor network
18:   $\mathcal{N} \leftarrow$ sample from normal distribution with mean 0 and variance $\sigma^2$
19:   $\mu_{noisy} \leftarrow \mu_{pred} + \mathcal{N}$ ▷ Inject exploration noise
20:   $\mu_{noisy}[w] \leftarrow \text{clamp}(\mu_{noisy}[w], 0, 1)$ ▷ Clamp wait fraction to be between 0 and 1
21:   $\mu_{noisy}[\Phi] \leftarrow \text{clamp}(\mu_{noisy}[\Phi], -\pi, \pi)$ ▷ Clamp heading angle to be between $-\pi$ and $\pi$

22:   // 3. Environment execution and evaluation
23:   $\mathbf{y}(T) \leftarrow$ simulate kinematics of the fish using $\mathbf{y}(0)$ and $\mu_{noisy}$
24:   $R \leftarrow \|\mathbf{x}(T) - \mathbf{y}(T)\|$ ▷ Calculate magnitude of interception error

25:   // 4. Critic network update (minimizing prediction error)
26:   $Q_{pred} \leftarrow Q(\mathbf{s}, \mu_{noisy} | \vartheta_Q)$ ▷ Forward execution through critic
27:   $\mathcal{L}_Q(\vartheta_Q) \leftarrow \sum (Q_{pred} + R)^2 / N$ ▷ Compute critic loss over batch of size $N$
28:   Compute gradient $\nabla_Q \mathcal{L}_Q(\vartheta_Q)$ via backpropagation
29:   Update critic parameters $\vartheta_Q$ using Adam and $\nabla_Q \mathcal{L}_Q(\vartheta_Q)$

30:   // 5. Actor network update (maximizing expected reward)
31:   $Q_{eval} \leftarrow Q(\mathbf{s}, \mu_{pred} | \vartheta_Q)$ ▷ Evaluate actions with updated critic parameters
32:   $\mathcal{L}_\mu(\vartheta_\mu) \leftarrow -\sum Q_{eval} / N$ ▷ Compute actor loss over batch of size $N$
33:   Compute gradient $\nabla_\mu \mathcal{L}_\mu(\vartheta_\mu)$ via backpropagation ▷ Chain rule through critic network
34:   Update actor parameters $\vartheta_\mu$ using Adam and $\nabla_\mu \mathcal{L}_\mu(\vartheta_\mu)$
35: end for

36: Return $\vartheta_\mu, \vartheta_Q$
```
